# Bibliometric analysis of neuroscience publications quantifies the impact of data sharing

**DOI:** 10.1101/2023.09.12.557386

**Authors:** Herve Emissah, Bengt Ljungquist, Giorgio A. Ascoli

## Abstract

**Motivation:** Neural morphology, the branching geometry of neurons and glia in the nervous system, is an essential cellular substrate of brain function and pathology. Despite the accelerating production of digital reconstructions of neural morphology in laboratories worldwide, the public accessibility of data remains a core issue in neuroscience. Deficiencies in the availability of existing data create redundancy of research efforts and prevent researchers from building on others’ work. Data sharing complements the development of computational resources and literature mining tools to accelerate scientific discovery.

**Results:** We carried out a comprehensive bibliometric analysis of neural morphology publications to quantify the impact of data sharing in the neuroscience community. Our findings demonstrate that sharing digital reconstructions of neural morphology via the NeuroMorpho.Org online repository leads to a significant increase of citations to the original article, thus directly benefiting the authors. Moreover, the rate of data reusage remains constant for at least 16 years after sharing (the whole period analyzed), altogether nearly doubling the peer-reviewed discoveries in the field. Furthermore, the recent availability of larger and more numerous datasets fostered integrative meta-analysis applications, which accrue on average twice the citations of re-analyses of individual datasets. We also designed and deployed an open-source citation tracking web-service that allows researchers to monitor reusage of their datasets in independent peer-reviewed reports. These results and the released tool can facilitate the recognition of shared data reuse for promotion and tenure considerations, merit evaluations, and funding decisions.

**Availability and implementation:** The code is available at https://github.com/HerveEmissah/nmo-authors-app (author app) and https://github.com/HerveEmissah/nmo-bibliometric-analysis (bibliometric analysis app).

## 1. Introduction

Omics and structural biology have benefited enormously from the consistent practice of data sharing, with thriving research subfields fueled by seminal discoveries entirely based on publicly available datasets, and vibrant ecosystems of related scientific tools (Field et al., 2009; Chervitz et al., 2011; Wilson et al., 2021). Neuroscience has followed suite only more recently and more gradually, in part due to the greater heterogeneity of data types and the lack of a clearly understood functional code akin to that of genomic sequences (Gardner et al., 2003; Gleeson et al., 2017; Poline et al., 2022). One particular domain of neuroscience, digital reconstructions of neural morphology, is especially amenable to data sharing (Ascoli, 2006; Ascoli, 2015; Ascoli, 2017).

The accelerating development of advanced technologies in microscopic imaging and computational processing has greatly enhanced the methods of 3D neural reconstruction, enabling the creation of ever larger amounts of digital tracing data (Ledderose et al., 2014; Liu et al., 2022; Manubens-Gil et al., 2023). Capitalizing on this growth requires effective data accessibility to accelerate scientific discovery in neuroscience. Indeed, this was the goal of NeuroMorpho.Org, an open-access archive of 3D neural reconstructions and associated metadata (Ascoli et al., 2007). Today, this resource comprises hundreds of thousands of downloadable reconstructions with their associated metadata, each of them linked to peer-reviewed publications from laboratories worldwide (Akram et al., 2018). Global collaborative efforts and data sharing from multiple sources are extremely valuable to researchers to gain a better understanding of the brain and its cellular constituents given the strong association between neuronal form and function (Parekh and Ascoli, 2015; Suárez et al., 2020). It is essential to determine, however, the effective extent and impact of free data exchange.

Previous research quantified the benefits of data sharing to the original authors who shared data, in addition to the data users and the community at large, in specific disciplines such as cancer microarray clinical trials (Piwowar, Day & Fridsma, 2007) and non-invasive human brain imaging (Milham et al, 2018). However, it is not yet known whether these findings generalize to other fields, and in particular if neural morphology data sharing provides a positive return on investment for the original data owners and/or significantly impacts scientific throughput.

We use computational and data retrieval techniques to quantitatively examine the patterns of publications, citations, and data reusage among researchers. Here we present a comprehensive bibliometric analysis of published literature pertaining to neural morphology to assess the impact of data sharing on the overall field as well as on individual investigators. We further introduce a dynamic web-based research tool to determine the scientific impact of uniquely identified, shared neural morphology datasets. The application serves as a valuable resource for neuroscientists to demonstrate the direct and indirect benefits of sharing their data.

## 2. Materials and methods

This study relies on datasets retrieved from the following repositories: NeuroMorpho.Org, Semantic Scholar, and Europe PubMed Central (EuropePMC). Semantic Scholar is an Artificial Intelligence-powered engine for research literature including a large neuroscience collection (Jones, 2015). Europe PMC is an open-access archive of life science publications (Ferguson et al., 2021). We have selected these databases due to their extensive full-text record coverage and accessibility via Application Program Interface (API). In particular, NeuroMorpho.Org tallies availability and reusage of neural morphology data, while Semantic Scholar and EuropePMC track peer-reviewed citations, broadly considered an expedient proxy for scientific impact.

We refer here to “primary” publications that generated new digital reconstructions of neural morphology as ***<u>Describing</u>***. NeuroMorpho.Org divides *Describing* publications into three categories depending on whether the underlying datasets are publicly available (***<u>Sharing</u>***), unavailable (***<u>Unsharing</u>***), or determining availability. The database curators determine this information through direct interaction with data owners (Maraver et al., 2019) and update it publicly every month (neuromorpho.org/LS_queryStatus.jsp?status=Available&page=0).

NeuroMorpho.Org also tracks the “secondary” publications that cite the *Describing* articles and/or utilize the corresponding downloaded digital reconstructions, referred to as ***<u>Citing</u>*** and ***<u>Using</u>***, respectively (neuromorpho.org/LS_usage.jsp). We fetch the *Describing, Citing* and *Using* metadata via the NeuroMorpho.Org API (neuromorpho.org/apiReference.html#literature) using a Python application, implemented with the Flask framework and released open source (https://github.com/HerveEmissah/nmo-bibliometric-analysis), to populate a MongoDB database. We retrieve the upload date as well for each dataset from the NeuroMorpho API (neuromorpho.org/api/neuron) and store it in the MongoDB database.

We also fetch citations to and references of primary and secondary publications programmatically for storage in the MongoDB database. Specifically, the Semantic Scholar API (api.semanticscholar.org/v1/paper/{doi}) returns a JSON formatted response containing both citations and references metadata for a given publication. EuropePMC, in contrast, exposes distinct API endpoints for citations (ebi.ac.uk/europepmc/webservices/rest/MED/{pmid}/citations) and references (ebi.ac.uk/europepmc/webservices/rest/MED/{pmid}/references). We use a union of citations and a union of references from both Semantic Scholar and EuropePMC to provide a more complete record.

The MongoDB database serves both as the repository for the bibliometric analysis (utilizing the plotting library Matplotlib for graphical visualization) and as the data source for a three-tier web-based author service, with a front-end user interface built with the React Framework and the back-end application developed in Python using Flask. The application components are built as separate Docker images exchanging information via a Docker network (Fig. 1). The bibliometric author service is deployed on a publicly accessible George Mason University server (cng-nmo-dev3.orc.gmu.edu:8181/) and is released open source as well (https://github.com/HerveEmissah/nmo-authors-app).

**Figure 1:**
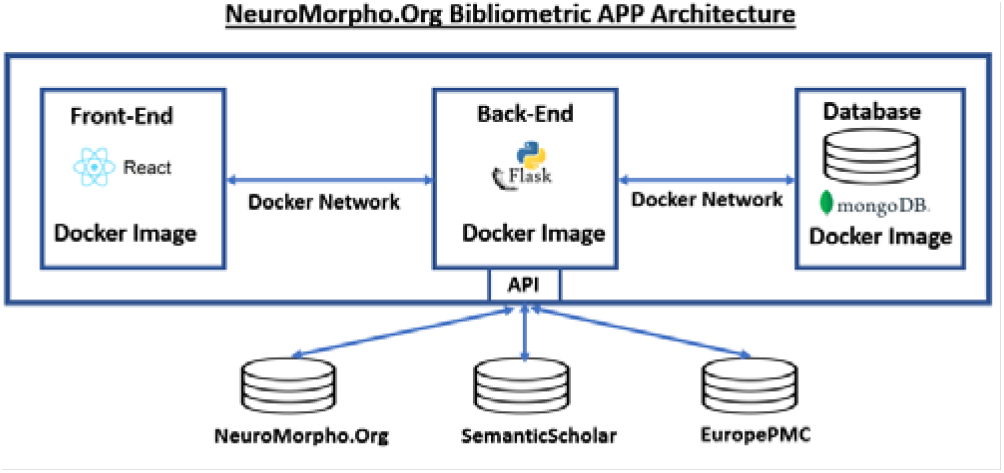
High-level schematic of the bibliometric author service 3-tier architecture, with a front-end presentation tier, Flask application, and MongoDb database.

## 3. Results

We first investigated whether openly sharing via NeuroMorpho.Org the digital reconstructions of neural morphology described in an article increases the number of citations to that article (Fig. 2). We started by comparing the number of citations to *Sharing* (N=1,656) and *Unsharing* (N=3,089) articles. Specifically, we normalized the yearly number of citations for a given *Describing* article by dividing its accrued citations by the number of years elapsed since publication. The analysis (Fig. 2A) demonstrates a significant difference in yearly citations between groups (*Sharing*:8.91 ± 14, *Unsharing*: 6.19 ± 12; effect size +43.9%, p=0.006).

**Figure 2.**
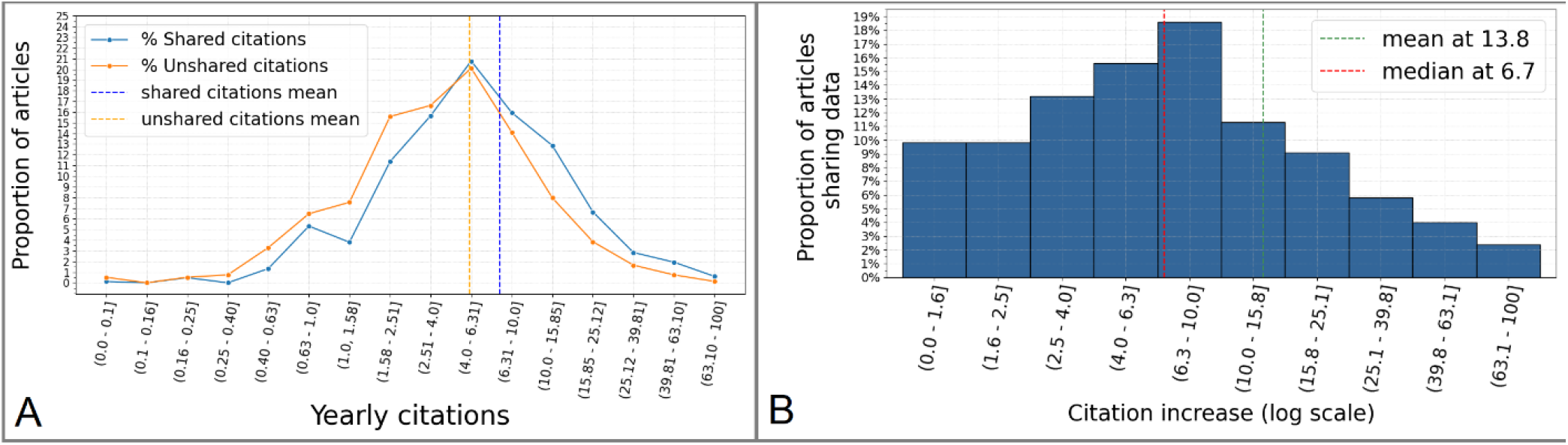
Publicly sharing digital reconstructions of neural morphology increases the number of citations to the *Describing* article. **A:** Distributions of citations for *Sharing* and *Unsharing* articles bin-grouped using logarithmic scale. **B:** Relative increase of citations to *Sharing* articles specifically due to *Citing* and *Using* publications.

We then asked whether this increase was specifically due to the citations by the *Citing* and *Using* publications. Thus, we calculated the **Citation Increase** for each *Sharing* article based on the following formula: **Citation Increase = NMO_Citations / (Citations_since_upload - NMO_Citations)**, where **NMO_Citations** represents the citations to the *Sharing* article by the *Citing/Using* publications, and **Citations_since_upload** represents the overall citations to the *Sharing* article since the upload date of the corresponding dataset. The resultant histogram distribution (Fig. 2B) reveals that the Citation Increase of *Sharing* articles due to the secondary publications (13.8%) explains less than a third of the difference in citations between *Sharing* and *Unsharing* articles. Taken together, these analyses indicate that sharing neural reconstruction data through NeuroMorpho.Org increases the impact of the original publication.

Next, we explored if the numbers of citations to *Sharing* articles due to secondary publications decreases over time after publication. The results suggest a broadly uniform citation likelihood without a tendency to decrease over the whole 16 years of the project activity (Fig. 3).

**Figure 3.**
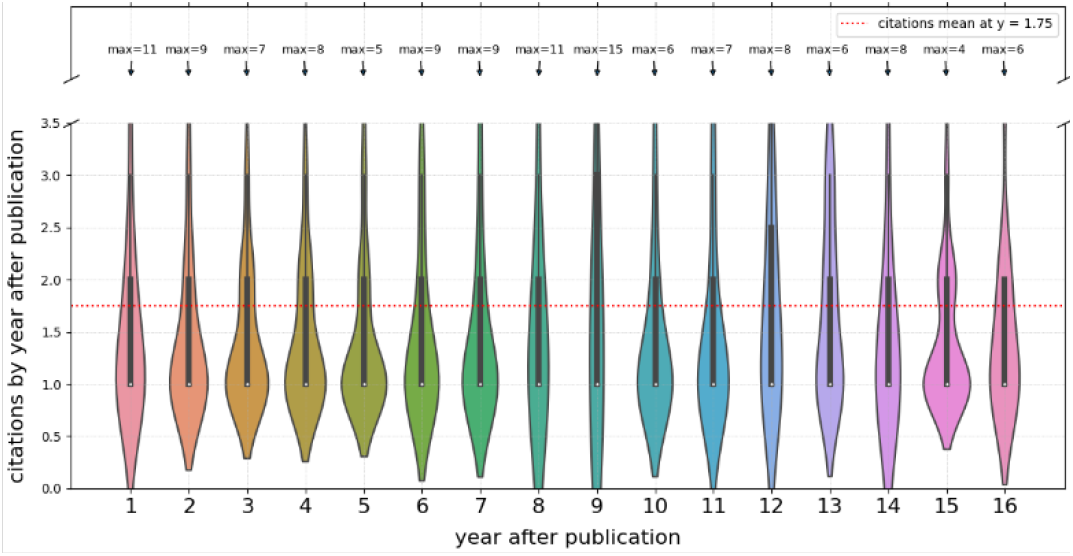
Yearly citations to *Sharing* article by *Citing* and *Using* publications as a function of the time elapsed since the publication of the *Sharing* article.

To help assess the impact of shared data, it is also interesting to compare the number of primary and secondary publications and their respective citations (Fig. 4). Both the cumulative number of *Describing* articles and of *Citing/Using* articles increased consistently from the project launch to present (Fig. 4A). Notably, the secondary publications, which rely on shared data, effectively double the primary (*Describing*) data literature. Moreover, comparing the overall citations to primary and secondary articles (Fig. 4B) again demonstrates that secondary publications increase the number of citations in the field by nearly 50%. These results further underscore the added impact of data sharing in neuroscience.

**Figure 4.**
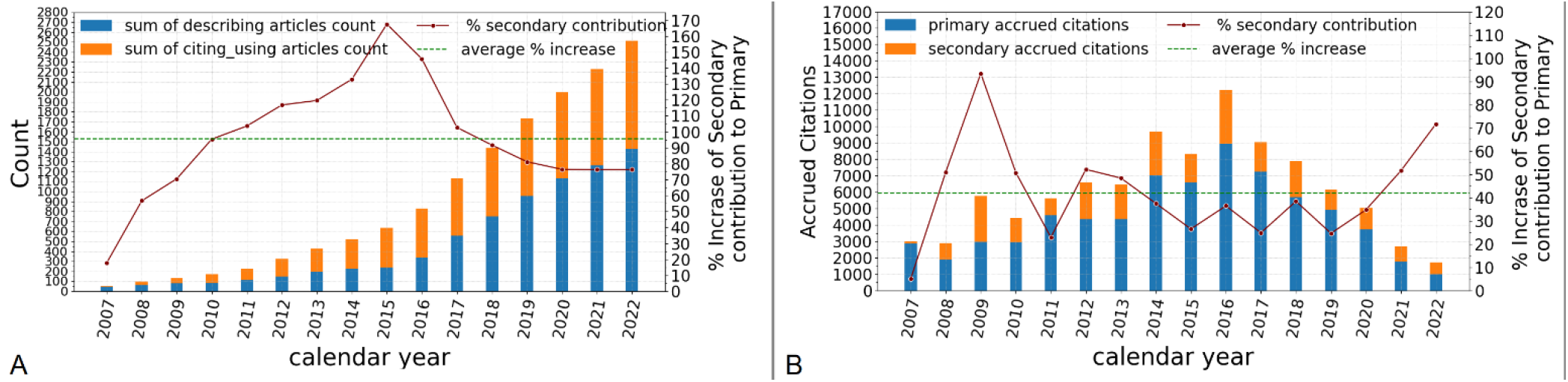
Impact of neural morphology data sharing on publications and citations. **A:** Cumulative sum of primary and secondary article counts by year. **B:** Citations accrued by primary and secondary articles by year.

We also analyzed how many primary articles are used or cited by secondary publications (Fig. 5). We found that fewer than a third of secondary studies only refer to a single source of primary data, and approximately the same proportion relies on two or three sources (Fig. 5A). In contrast, more than 40% of secondary articles cite or use 4 or more sources, and nearly 10% require ten or more primary publications. Interestingly, the ratio between the number of secondary publications citing or using 4 or more primary articles and the number of secondary publications citing or using 3 or fewer primary articles (Fig. 5A inset) is substantially greater in the scientific literature after 2016 (ratio: 0.82) than before (ratio: 0.54), reflecting the increasing emphasis on big science, data aggregation, and meta-analyses, while previous studies may have been concerned with case-by-case morphology.

**Figure 5.**
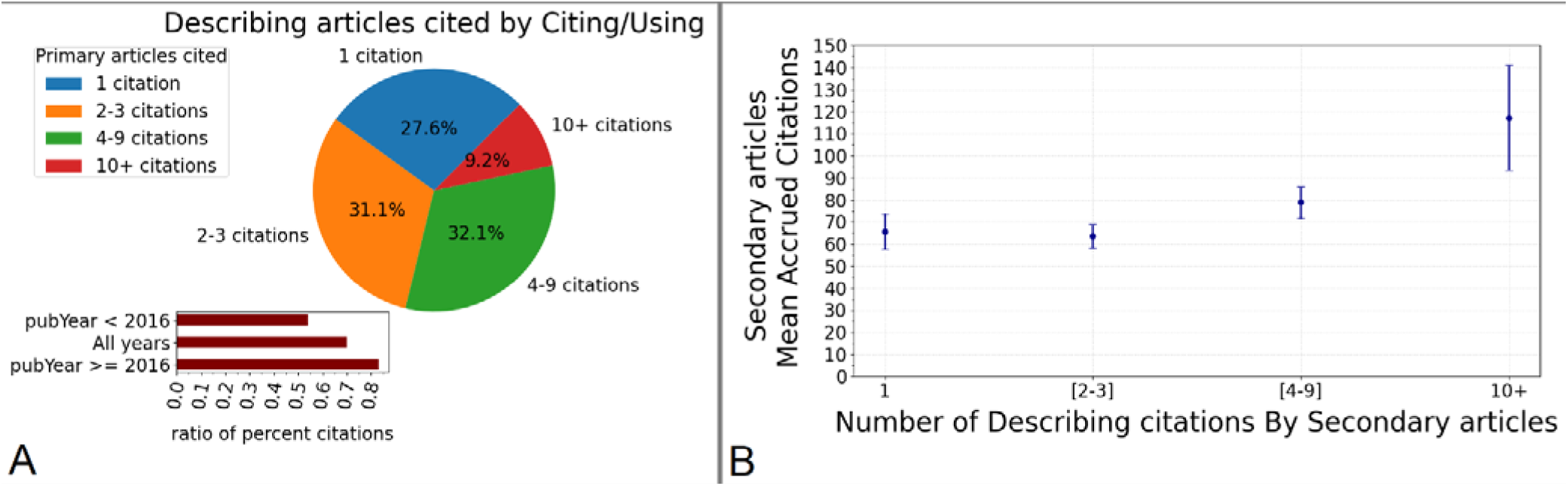
Integration of multiple data sources in secondary publications. **A:** Proportion of secondary publications citing or using different numbers of primary articles. **Inset:** ratio between the number of secondary publications citing or using 4 or more primary articles and those citing or using 3 or fewer primary articles. **B:** Mean number of citations accrued by secondary publications as a function of the number of *Describing* articles cited. The error bars indicate 95% confidence interval.

Furthermore, we asked whether the secondary articles using a greater number of primary sources are cited more than those using fewer primary sources (Fig. 5B). Indeed, while secondary publications using 1-3 primary sources only accrue on average 60-65 citations, the mean number of accrued citations reaches approximately 80 for secondary articles using 4-9 sources and exceeds 110 for secondary articles using 10 or more primary sources.

To provide researchers the capability to investigate the impact of *Sharing* articles on secondary publications, we made the bibliometric functionality utilized in the above analysis available as a public service through a web-based user interface (Fig. 6). The service fetches the list of *Sharing* publications from the NeuroMorpho.Org datastore and renders the corresponding unique identifiers (PMID/DOI) on the application interface, allowing the user to select interactively the publication(s) of interest for the retrieval of secondary dataset citations (Fig. 6A). The Get Alert function enables subscribing to automated email notifications when the dataset from an article of interest is cited or reused (Fig. 6B) and allows users to unsubscribe whenever desired.

**Figure 6:**
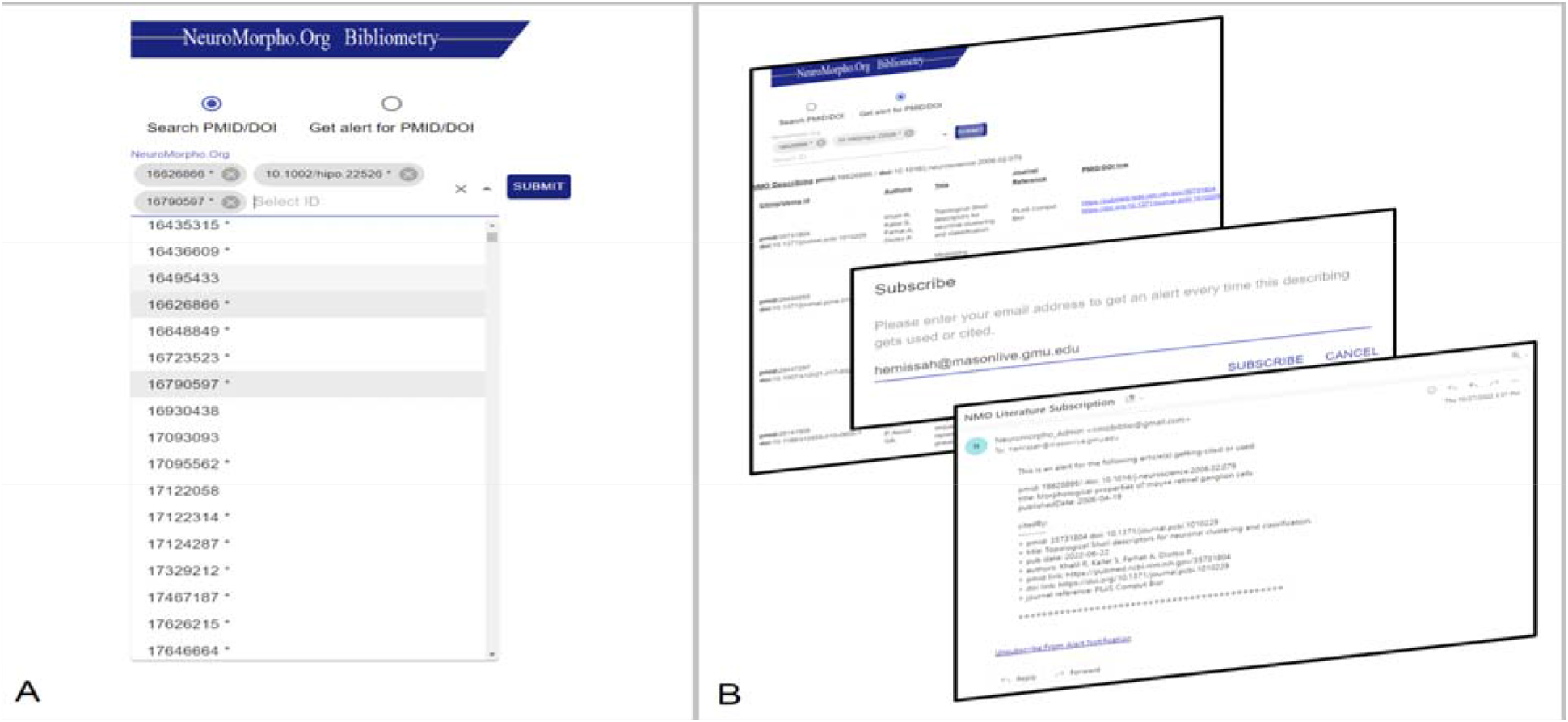
Bibliometric Author Service. **A**. The user interface lists all *Describing* publications, allowing one or more to be selected for analysis of secondary usage or subscription to alert notification. **B**. Display of secondary usage provides bibliographic details of the secondary publications pertaining to the selected article. Additionally, when subscribing to citation alerts for specific publications, users receive email notifications when datasets from their publications of interest are reused or cited.

## 4. Discussion

The analysis of neuroscience data sharing practice among researchers provides insight into current trends and raises awareness to encourage more collaborations and open release of data (Poldrack & Gorgolewski, 2014). It is intuitively obvious that the public availability of data is beneficial to researchers who can reuse it for secondary analysis and diverse scientific applications (Halavi et al., 2012). As neural morphology datasets are increasing in scale, it is important to enable broad community access. This can be facilitated by encouraging authors to consistently share their data. However, whether it in fact provides advantages for the data owners to share data has remained a topic for discussion.

Our findings demonstrate indeed that sharing digital reconstructions of neural morphology via the NeuroMorpho.Org online repository leads to a significant increase of citations to the original article, thus directly benefiting the authors. Moreover, the rate of data reusage remains constant for at least 16 years after sharing (the whole period analyzed), altogether nearly doubling the peer-reviewed discoveries in the field. Furthermore, the recent availability of larger and more numerous datasets fostered integrative meta-analysis applications, which accrue on average twice the citations of re-analyses of individual datasets. These results demonstrate the broader impact of open sharing of neural reconstructions on scientific discovery.

We also designed and deployed an open-source bibliometric tracking web-service that allows researchers to monitor reusage of their datasets in independent peer-reviewed reports. This computational tool can facilitate the recognition of shared data reuse for promotion and tenure considerations, merit evaluations, and funding decisions.

## Acknowledgments

This research was supported in part by NIH grants R01NS39600 and R01NS86082. The authors are grateful to Dr. Patricia Maraver for guidance and help with the usage of the NeuroMorpho.Org literature management system.

## Notes

### Competing Interest Statement

The authors have declared no competing interest.

